# Clade GR and Clade GH Isolates in Asia Show Highest Amount of SNPs

**DOI:** 10.1101/2020.11.30.402487

**Authors:** Antara Sengupta, Sk. Sarif Hassan, Pabitra Pal Choudhury

## Abstract

Clades are monophyletic groups composed of a common ancestor and all its lineal descendants. As the propensity of virulence of a disease depends upon the type of clade the virus belongs to and it causes different fatality rates of disease in different countries, so the clade-wise analysis of SARS-CoV-2 isolates collected from different countries can illuminate the actual evolutionary relationships between them. In this study, 1566 SARS-CoV-2 genome sequences across ten Asian countries are collected, clustered, and characterized based on the clade they belong to. The isolates are compared to the Wuhan reference sequence (Accession no:*MN*996528.1) to identify the mutations that occurred at different protein regions. Structural changes in amino acids due to mutations lead to functional instability of the proteins. Detailed clade-wise functional assessments are carried out to quantify the stability and vulnerability of the mutations occurring in SARS-CoV-2 genomes which can shade light on personalized prevention and treatment of the disease and encourage towards the invention of clade-specific vaccines.

## 1. Introduction

Viruses have a remarkable capacity to adapt to new hosts and environments [1]. Mutations may lead to different phenotypic changes in them, which may lead to occur biodiversity. Phylogenies are frameworks for analysing biodiversity. Phylogenetic analysis based on sequence similarity is one of the very efficient way to do so [2]. However, it will be worth noting that due to the recent outbreak of pandemic COVID-19, people around the world are trying by every means to reach the origin, to get some ways of prevention and therapeutic pathways. Biodiversity is characterized by a continual replacement of branches in the tree of life, i.e. clade [3]. Evolutionary pressure on host immunodeficiency leads to different clades of viruses [4]. A clade is a group of highly related sequences that share a common ancestor. They can provide hypotheses about the actual evolutionary history of that group of sequences. Some clinical studies suggest that the proclivity of virulence of a disease depends upon the type of clade the virus belongs to [4]. Clade differences can result in varying degrees of pathology. Millions of gene regulatory elements are there which contribute heavily to the variation in gene expression of complex human traits and diseases [5].Determining mutation types influence a lot in gene regulation and is important for studying the role of regulatory variation in evolution. Genomic evolution helps a virus to escape host immunity [6, 7]. The clade-wise analysis of SARS-CoV-2 isolates collected from different countries can shed a light on the actual evolutionary history of the region or continent. In order to confirm the hypothesis in COVID-19 pathogenesis, it is highly recommended to make a thorough study of mutations occurring in SARS-CoV-2 isolates collected from different demographic areas and characterizing them based on the clades they come from [8]. A plethora of papers already have been published, where researchers have tried to study the virus isolates of SARS-CoV-2, which is solely responsible for the disease to occur in human [9, 10, 11, 12, 13]. Huge numbers of investigations are reported in order to find evolutionary relationships between SARS-CoV-2 and other corona-viruses and to determine the origin and molecular characteristics of SARS-CoV-2 [14, 15, 16]. Several works are also done on characterization and comparative analysis of structured and non-structured proteins of SARS-CoV-2 [17, 18]. According to Hassan et.al. [19] among all the accessory proteins of SARS-CoV-2, ORF3a plays an important role in virus pathogenesis, as it possesses various mutations which are linked with that of spike proteins.It is observed that due to its structural plasticity and high diversity, ORF8 plays important role in SARS-CoV-2 pathogenicity [20].Computational biology approaches are applied to investigate genomic and proteomic variations of S protein in SARS-CoV-2. In the paper authors find possibilities to design potential inhibitors against S protein [21]. Kumar et.al.stated the first observation on the deletion mutations in the C-terminal region of the envelope glycoprotein in India [22]. Some researchers have given more stresses on codon usage bias in SARS-CoV-2 rather than mutational trends [23]. Several investigations are carried out on characterizing the mutations of a particular country and even across the globe [24, 25, 26]. Researchers aimed too to analyse SARS-CoV-2 proteins modulating host immune response like type I interferon pathways [27]. People have tried to uncover the relation between hotspot mutations and viral pathogenicity [28]. Some research papers focused on characterizing B and T cell epitopes of certain proteins of SARS-CoV-2 which can help in vaccine development [29]. Phylodynamic analyses of SARS-CoV-2 genomes can provide insights into the roles of some relevant factors to limit the spread of the disease [30]. Priya and Shanker [31] have observed that the coevolutionary forces can increase the fitness of spike glycoprotein against ACE2 and increase the infectivity of SARS-CoV-2. According to Uddin et al. [32] antigenic epitopes of SARS-CoV-2 and SARS-CoV have highest level of similarities among them.SARS-CoV-2 is the seventh coronavirus to infect humans but the first HCoV which pandemic potential [33]. (Accession no:*NC*_045512) is the first SARS-CoV-2 sample SARS-CoV-2 sequence from Wuhan, and it is from clade ’O’ [34]. Clade ’G’ is the variant of the spike protein D614G which indicates significantly higher human host infectivity and better transmission efficiency to the virus. GH and GR are the most common offsprings of clade G. According to data from the public database of the Global Initiative on Sharing All Influenza Data (GISAID), three major clades of SARS-CoV-2 are clade G (variant of the spike protein S-D614G), clade V (a variant of the ORF3a coding protein NS3-G251), and clade S (variant ORF8-L84S) [34]. GR clade, carrying the combination of NSP3: F106F, Spike: D614G and Nucleocapsid: RG203KR mutations, whereas clade ’GH’ represents the mutations NSP12b: P314L, S: D614G and ORF3a: Q57H. Different fatality rates observed in different countries may be the consequence of clade’s differences in virulence. The spike protein of SARS-CoV-2 binds the host receptor angiotensin-converting enzyme 2 (ACE2) via receptor-binding domain (RBD). It is reported that immunization with SARS-CoV-2 receptor-binding domain (RBD) is able to induce clade-specific neutralizing antibodies in a host like mice [35]. In some cases vaccines are immunogenic and induced antibodies can neutralize homologous and heterogeneous viruses with different degrees of cross-reactivity [36]. Hence, in this present study, 1566 SARS-CoV-2 isolates from the Asian continent comprising 10 countries (India, Bangladesh, Pakistan, Srilanka, China, Japan, Malaysia, Iran, Thailand, and Saudi Arabia) are collected, clustered, and characterized based on the clade they belong to.

## 2. Methods and Materials

### 2.1. Collection of gene sequences of SARS-COV-2

One of the primary features of the investigation and analysis of the COVID-19 is availability of real-time data in global databases. To carry out the experiment We have collected 1566 isolates of SARS-CoV-2 from ten different Asian countries from the National Center for Biotechnology Information (NCBI) database (https://www.nih.gov/coronavirus) on October 20, 2020. The information about collected dataset are presented as Supplemental Materials in Table S1 and summarized in Table 1. In addition, we have collected the Reference Sequence (Accession no:*MN*996528.1) from the same Gene bank.However, information in detail about the dataset Collected sequences are then gone through preliminary screening for excluding noisy sequences. Here noise includes no mutations and the amino acid changes due to mutations specified by ’X’. Thus finally 1384 isolates are taken for further investigations.

**Table 1:** Collected SARS-CoV-2 genome sequences from COVID patients from ten different Asian Countries.

| Country Name | # Associated Isolate | #Isolate in work |
| --- | --- | --- |
| INDIA | 570 | 565 |
| THAILAND | 227 | 104 |
| BANGLADESH | 231 | 231 |
| IRAN | 172 | 106 |
| CHINA | 189 | 189 |
| JAPAN | 96 | 96 |
| SAUDI ARABIA | 58 | 58 |
| PAKISTAN | 10 | 9 |
| MALAYSIA | 9 | 9 |
| SRILANKA | 4 | 4 |

### 2.2. Methods

The present work aims to make a clade-wise classification and analysis of SARS-CoV-2 isolates of ten Asian countries. The isolates of each country are then compared with the reference sequence to find out the mutations that occurred. Clade-wise clustering of the given dataset is taken place. The observed mutations are then gone through different online software tools to investigate different biological functionalities that may change and affect the variants due to mutations. Here it is to be noted that we have used two web-based software tools (PROVEAN [37] and I-mutant [38]) for the aforesaid functional assessments.I-Mutant is a suite developed based on Support Vector Machine(SVM). (ΔΔ*G*>−0.5 Kcal/mol) indicates that the mutation can largely destabilize the protein, ΔΔ*G* >0.5 Kcal/mol indicates about the strong stability and −0.5>=ΔΔ*G*>=0.5 Kcal/mol tells about weak effect of mutations. Isolates with a score equal to or below −2.5 are considered deleterious and scores above −2.5 are neutral. Lastly, we tracked the trend of mutations that occurred in the sequences of different clades.

## 3. Results and Discussions

### 3.0.1. Clade-wise clustering of SARS-CoV-2 strains taken as dataset from different countries

After excluding the noisy sequences finally 1371 isolates are found. Each strain belongs to a particular clade, so the isolates are clustered according to the clade from which they belong to. It has been observed that as a whole isolates of five clades (G, GH, GR, L, S, O, and V) are participated in those countries of the Asian continent. According to (Fig. 1) the order of the clade-wise participation of isolates is GH>GR>O>G>S>L>V. It is to be noted here that among the entire dataset taken Indian isolates hold a big amount of data. According to the country-wise view shown in Table 2 SARS-CoV-2 isolates of clade ’O’ are present in the dataset of all countries and isolates of clade ’V’ have been circulated only at China and Thailand. The country-wise analysis has a mixed result. In Srilanka,Thailand, China, Malaysia, and Iran the isolates are majorly from clade ’O’. India and Saudi Arabia have a prevalence of clade ’GH’. Pakistan, Bangladesh, and Japan have the prevalence of clade ’GR’. It indicates viral diversity regarding infection as the infection is transmitting from one country to another. Remarkable viral diversities are also present even in different regions within a country too.

**Figure 1:**
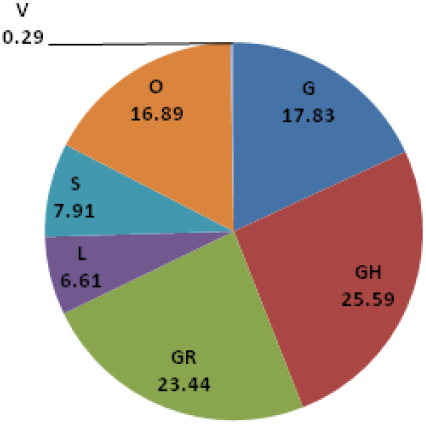
Calculate percentage of clades representing the isolates.

**Table 2:** Clade-wise counting of isolates reported in 10 countries.

| COUNTRY | G | GH | GR | L | S | O | V |
| --- | --- | --- | --- | --- | --- | --- | --- |
| THAILAND | 12 | 0 | 0 | 1 | 39 | 51 | 1 |
| BANGLADESH | 6 | 13 | 208 | 1 | 1 | 2 | 0 |
| SRILANKA | 1 | 0 | 1 | 0 | 0 | 2 | 0 |
| CHINA | 2 | 0 | 1 | 64 | 48 | 71 | 3 |
| JAPAN | 2 | 5 | 71 | 13 | 2 | 3 | 0 |
| INDIA | 191 | 303 | 18 | 7 | 19 | 27 | 0 |
| IRAN | 32 | 1 | 0 | 4 | 0 | 69 | 0 |
| MALAYSIA | 0 | 1 | 0 | 0 | 0 | 8 | 0 |
| PAKISTAN | 0 | 1 | 5 | 2 | 0 | 1 | 0 |
| SAUDI ARABIA | 2 | 32 | 22 | 0 | 1 | 1 | 0 |

### 3.1. Investigating trend of mutations in various clades

In this subsection firstly the positions of mutations are identified in each isolate and then it is aimed to calculate clade-wise percentage of mutations occurred in each country as shown in (Fig. 2).

**Figure 2:**
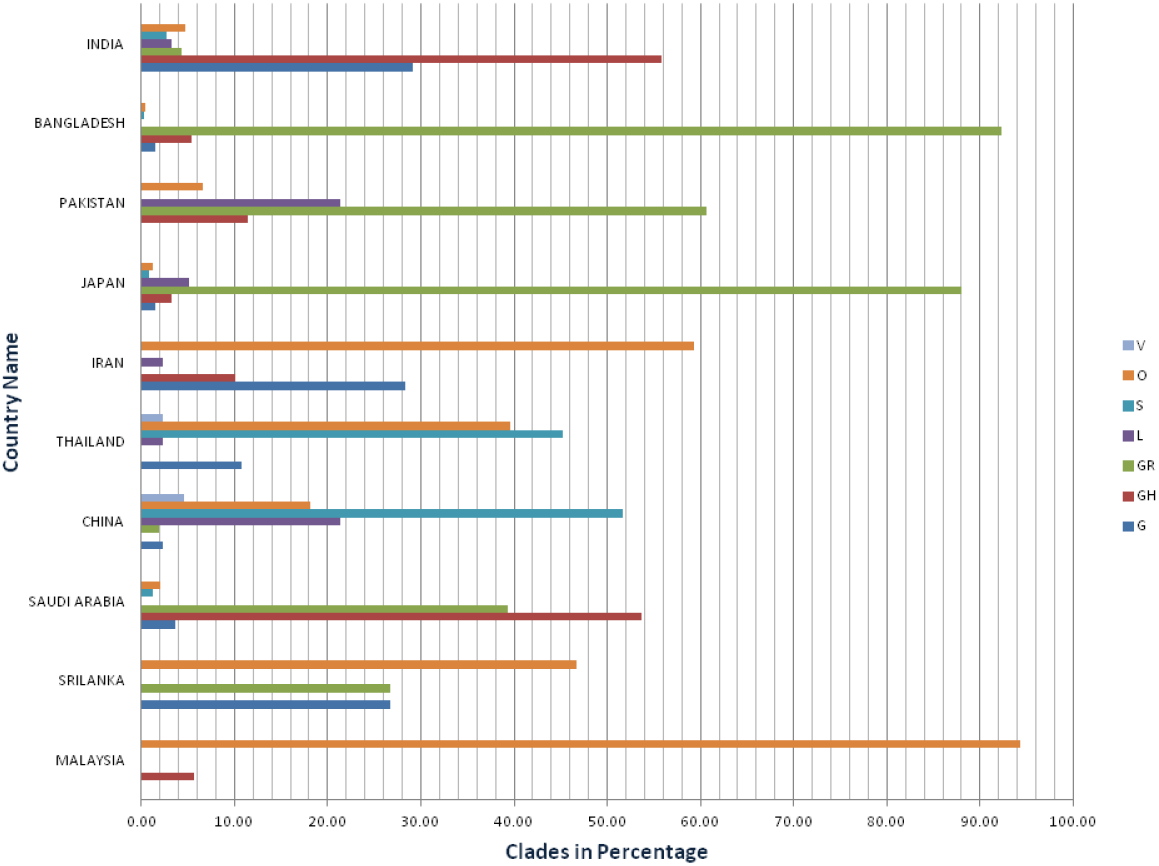
Calculating clade-wise percentage of mutations occurred in different countries.

Secondly, a microscopic view has been given on clade-wise clustering of total mutations found in the whole dataset and calculating the protein-wise percentage of the mutations occurred according to the clades they belong to (Fig. 3).

**Figure 3:**
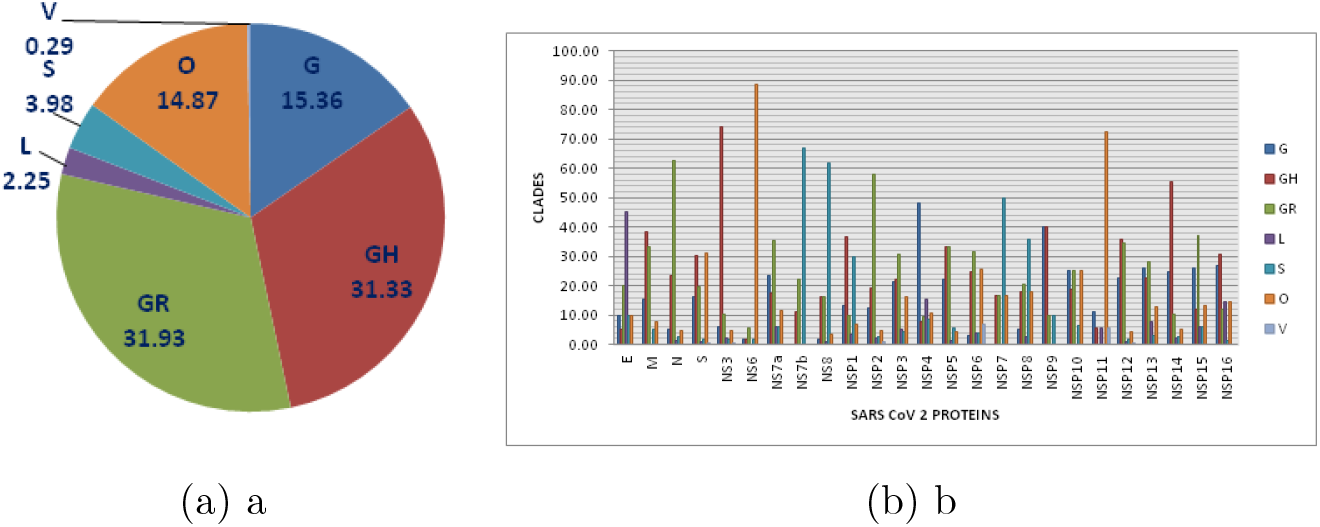
Quantitative analysis of clade-wise non-synonymous mutation. (a) Percentage of non-synonymous mutations found in different clades; (b) Clade-wise percentage of non-synonymous mutations occurred in different protein regions

According to (Fig. 2), India and Saudi Arabia have a prevalence of clade ’GH’ (55.76%, and 53.69% respectively). Whereas, in Pakistan, Bangladesh, and Japan strains have a prevalence of clade ’GR’ (60.66%, 92.33% and 88.06% respectively). Mutations have occurred in strains from Clade ’S’ at China and Thailand (51.63% and 45% respectively). SARS-CoV-2 isolates of clade ’O’ have significant participations in Iran, Srilanka and Malaysia (59.31%, 46.67% and 94.29% respectively). SARS-CoV-2 isolates of clades G, GH, GR, S, L, and O are circulating in India and Japan. Whereas, clades V, O, S, L, GR, and G are circulating at different regions of China and clades O, S, GR, GH, and G are circulated in Saudi Arabia. In Malaysia (clades O and GH) and Srilanka (clades O, GR, and G) the SARS-CoV-2 isolates do not have the viral diversities a lot.

Mutations refer to the virus to undergo certain changes which can lead to develop some new isolates after replications. Non-synonymous substitutions play a very significant role as this type of mutation makes change in amino acid. Alteration in amino acid causes structural change. With the aim of understanding the trend of non-synonymous mutations in different clades in the context of disease severity, a detailed protein-wise comparative analysis has been taken place. Mutations identified at different protein regions in all the isolates are shown at Table S2 in Supplementary file. To do so we have considered the total dataset as a whole. Clade-wise percentages of non-synonymous mutations at different protein regions are calculated. The clade-wise characterization of mutations of different proteins are shown in (Fig. 3).

According to the dataset taken, we have got 6665 numbers of non-synonymous mutations. We can observe at Table 3 that the chronological order of clades at per number of mutations taken place in whole dataset is GR>GH>G>O>S>L>V. It can be observed in (Fig. 3) that mutations are majorly taken place at isolates of clades GH and GR which are 31.33% and 31.93% of respectively. Samples of clade V have been affected rarely (0.29%). Clade-wise distribution of mutations in each protein does not have a very similar trend(s). Although the majority of proteins mutated are either of clade GR (N, NS7a, NSP2, NSP6, NSP13, and NSP15) or GH (M, NS3, NSP1, NSP12, NSP14, and NSP16) but clade G also has large numbers of mutations in some proteins(S, NS6, and NSP11). Isolates of Clade L and clade G here have got mutations maximum only in protein E and NSP4 respectively. In proteins NS7b, NS8, and NSP7 of the isolates from clade S have maximum distributions of non-synonymous mutations. In the isolates of clade G, GR, and O number of mutations at NSP10 are equal. Mutations are also equally distributed in protein NSP5 of clades GH and GR.

**Table 3:** Clade-wise segregation of mutated data

| PROTEIN | G | GH | GR | L | S | O | V |
| --- | --- | --- | --- | --- | --- | --- | --- |
| E | 2 | 1 | 4 | 9 | 2 | 2 | 0 |
| M | 6 | 15 | 13 | 0 | 2 | 3 | 0 |
| N | 60 | 264 | 709 | 13 | 31 | 52 | 0 |
| NS3 | 30 | 376 | 709 | 12 | 10 | 24 | 3 |
| NS6 | 1 | 1 | 3 | 0 | 1 | 47 | 0 |
| NS7a | 4 | 3 |  | 1 | 1 | 2 | 0 |
| NS7b | 0 | 1 |  | 0 | 6 | 0 | 0 |
| NS8 | 2 | 18 | 18 | 1 | 69 | 4 | 0 |
| NSP1 | 4 | 11 |  | 1 | 9 | 2 | 0 |
| NSP10 | 4 | 3 | 4 | 0 | 1 | 4 | 0 |
| NSP11 | 2 | 1 |  | 1 | 0 | 13 | 1 |
| NSP12 | 241 | 383 | 366 | 9 | 18 | 45 | 3 |
| NSP13 | 24 | 21 | 26 | 7 | 3 | 12 | 0 |
| NSP14 | 49 | 109 | 20 | 4 | 5 | 10 | 0 |
| NSP15 | 22 | 10 | 31 | 5 | 5 | 11 | 0 |
| NSP16 | 20 | 23 | 9 | 11 | 1 | 11 | 0 |
| NSP2 | 52 | 82 | 245 | 9 | 11 | 20 | 4 |
| NSP3 | 109 | 114 | 156 | 27 | 21 | 82 | 0 |
| NSP4 | 50 | 8 | 10 | 16 | 9 | 11 | 0 |
| NSP5 | 16 | 24 | 24 | 1 | 4 | 3 | 0 |
| NSP6 | 3 | 25 | 32 | 4 | 4 | 26 | 7 |
| NSP7 | 0 | 1 | 1 | 0 | 3 | 1 | 0 |
| NSP8 | 2 | 7 | 8 | 1 | 14 | 7 | 0 |
| NSP9 | 4 | 4 | 1 | 0 | 1 | 0 | 0 |
| S | 317 | 583 | 384 | 18 | 34 | 599 | 1 |
| **TOTAL MUTATION** | **1024** | **2088** | **2128** | **150** | **265** | **991** | **19** |

### 3.2. Quantitative assessment of functional changes occur due to mutations

Structural changes in amino acids due to mutations lead to create functional instability of the isolates themselves, cause vulnerable diseases and even increase the magnitude of virulence. In this subsection, we have tried to find the impact of single point mutations on the biological function of proteins of each isolates through the light of PROVEAN (Protein Variation Effect Analyzer) score, which may be deleterious or neutral [39]. We have also calculated the change in Gibbs free energy (ΔΔ*G*) occur due to single point mutations as the difference in folding free energy change between wild type and mutant protein (ΔΔ*G*) is considered as an impact factor of protein stability changes [40]. The motivation here is to understand the effect of those mutations on protein stability. The quantitative analysis will give an insight into the probable mutations that occur in a particular clade and the magnitude of virulence of them. It is to be noted that here we have excluded the mutations which are occurred only once.The deleterious mutations are shown in Table 4. It is observed at Table 4 that if we consider the dataset as a whole, then among structural proteins the mutations occurred in spike protein(s) are more deleterious than others. Among the accessory proteins, NS3 is affected the most. NSP2, NSP5, and NSP12 are the non-structural proteins that have most of the deleterious mutations that occurred. Furthermore, we have calculated clade-wise percentage of deleterious mutations that occurred in different protein regions. To do so, we have segregated each deleterious mutation occurred in ten different countries along with their clades Table 5. The (Fig. 4) shows the protein regions that are mostly affected by the deleterious mutations. Maximum deleterious mutations occurred in structural and accessory proteins belong to clade GH. Most of the deleterious mutations in non-structural protein regions are occurred in the isolates of both clades GH and GR. The isolates of clade V are rare and only found in the isolates of China and Thailand, but interestingly it is observed that few of deleterious mutations are also enlisted there. (Fig.4) depicts the fact that most of the deleterious mutations take place in amino acid sequences of clade GH. It is reported that the human genome may carry large numbers of deleterious mutations which as a whole make a significant contribution to fatal diseases. Identification and analysis of deleterious mutations can shade lights on personalized treatment and medicine [41]. Hence, the identification of these kinds of mutations in SARS-CoV-2 isolates and their impacts on the host body seek attention of virologists.

**Figure 4:**
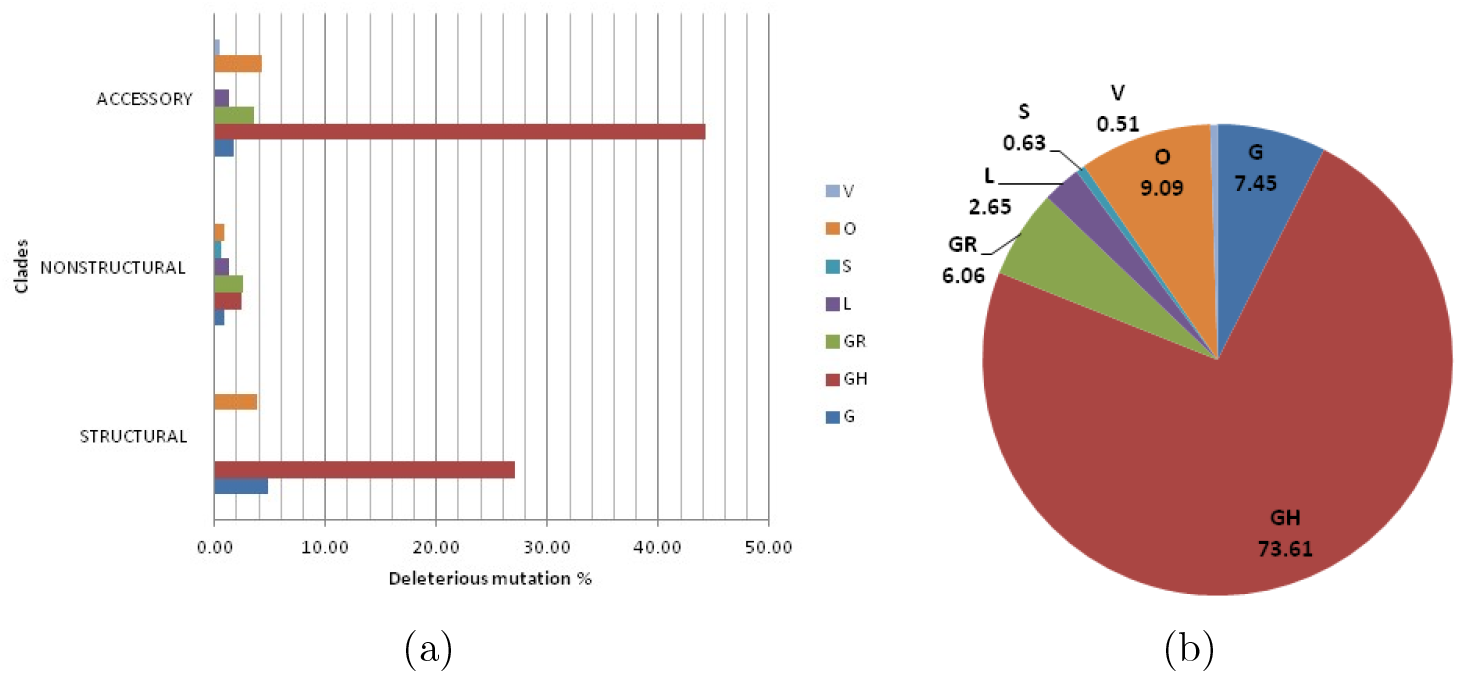
Quantification of clade-wise deleterious non-synonymous mutation. (a) Clade-wise percentage of deleterious non-synonymous mutations in different protein regions; (b) Clade-wise calculations of deleterious non-synonymous mutations of total data set

**Table 4:**
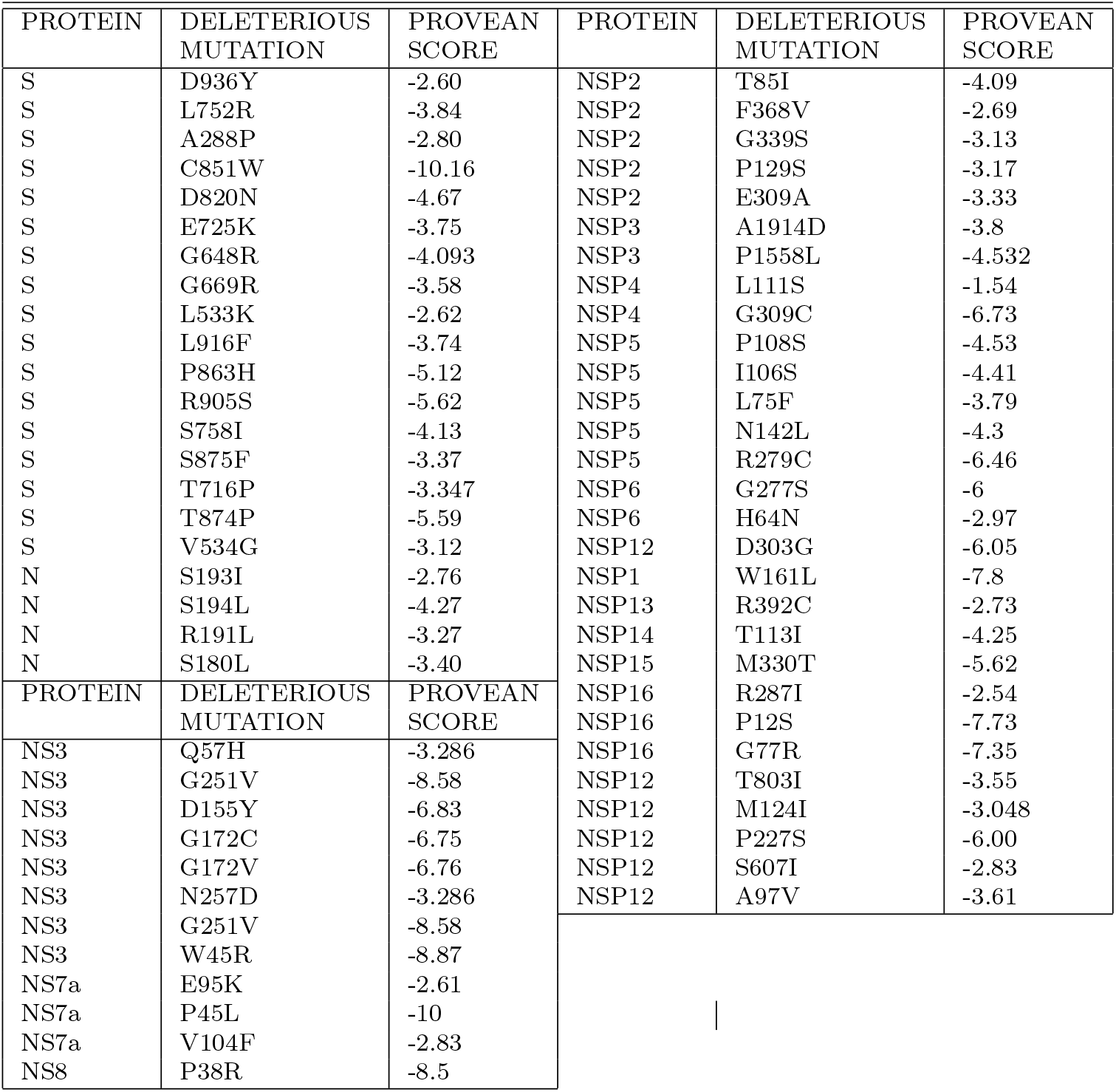
Investigate the deleterious mutations in total dataset.

| PROTEIN | DELETERIOUS MUTATION | PROVEAN SCORE | PROTEIN | DELETERIOUS MUTATION | PROVEAN SCORE |
| --- | --- | --- | --- | --- | --- |
| S | D936Y | -2.60 | NSP2 | T85I | -4.09 |
| S | L752R | -3.84 | NSP2 | F368V | -2.69 |
| S | A288P | -2.80 | NSP2 | G339S | -3.13 |
| S | C851W | -10.16 | NSP2 | P129S | -3.17 |
| S | D820N | -4.67 | NSP2 | E309A | -3.33 |
| S | E725K | -3.75 | NSP3 | A1914D | -3.8 |
| S | G648R | -4.093 | NSP3 | P1558L | -4.532 |
| S | G669R | -3.58 | NSP4 | L111S | -1.54 |
| S | L533K | -2.62 | NSP4 | G309C | -6.73 |
| S | L916F | -3.74 | NSP5 | P108S | -4.53 |
| S | P863H | -5.12 | NSP5 | I106S | -4.41 |
| S | R905S | -5.62 | NSP5 | L75F | -3.79 |
| S | S758I | -4.13 | NSP5 | N142L | -4.3 |
| S | S875F | -3.37 | NSP5 | R279C | -6.46 |
| S | T716P | -3.347 | NSP6 | G277S | -6 |
| S | T874P | -5.59 | NSP6 | H64N | -2.97 |
| S | V534G | -3.12 | NSP12 | D303G | -6.05 |
| N | S193I | -2.76 | NSP1 | W161L | -7.8 |
| N | S194L | -4.27 | NSP13 | R392C | -2.73 |
| N | R191L | -3.27 | NSP14 | T113I | -4.25 |
| N | S180L | -3.40 | NSP15 | M330T | -5.62 |
| PROTEIN | DELETERIOUS MUTATION | PROVEAN SCORE | NSP16 | R287I | -2.54 |
| NS3 | Q57H | -3.286 | NSP16 | P12S | -7.73 |
| NS3 | G251V | -8.58 | NSP16 | G77R | -7.35 |
| NS3 | D155Y | -6.83 | NSP12 | T803I | -3.55 |
| NS3 | G172C | -6.75 | NSP12 | M124I | -3.048 |
| NS3 | G172V | -6.76 | NSP12 | P227S | -6.00 |
| NS3 | N257D | -3.286 | NSP12 | S607I | -2.83 |
| NS3 | G251V | -8.58 | NSP12 | A97V | -3.61 |
| NS3 | W45R | -8.87 |  |  |  |
| NS7a | E95K | -2.61 |  |  |  |
| NS7a | P45L | -10 |  |  |  |
| NS7a | V104F | -2.83 |  |  |  |
| NS8 | P38R | -8.5 |  |  |  |

**Table 5:** Clade-wise clustering of deleterious mutations in total dataset.

| MUTATION | G | GH | GR | L | S | O | V | MUTATION | G | GH | GR | L | S | O | V |
| --- | --- | --- | --- | --- | --- | --- | --- | --- | --- | --- | --- | --- | --- | --- | --- |
| S193I | 1 | 7 | 0 | 0 | 0 | 0 | 0 | V104F | 0 | 2 | 0 | 0 | 0 | 0 | 0 |
| S194L | 21 | 190 | 0 | 0 | 0 | 2 | 0 | P38R | 0 | 0 | 3 | 0 | 0 | 0 | 0 |
| R191L | 2 | 0 | 0 | 0 | 0 | 0 | 0 | W161L | 1 | 1 | 0 | 0 | 0 | 0 | 0 |
| S180L | 1 | 1 | 0 | 0 | 0 | 0 | 0 | R392C | 0 | 0 | 2 | 0 | 0 | 0 | 0 |
| D936Y | 1 | 1 | 0 | 0 | 0 | 0 | 0 | T113I | 0 | 0 | 0 | 1 | 0 | 4 | 0 |
| L752R | 0 | 0 | 0 | 0 | 0 | 2 | 0 | M330T | 0 | 0 | 2 | 0 | 0 | 0 | 0 |
| A288P | 0 | 0 | 0 | 0 | 0 | 3 | 0 | R287I | 0 | 0 | 3 | 0 | 0 | 0 | 0 |
| C851W | 1 | 0 | 0 | 0 | 0 | 1 | 0 | P12S | 0 | 0 | 0 | 4 | 0 | 0 | 0 |
| D820N | 0 | 0 | 0 | 0 | 0 | 2 | 0 | G77R | 0 | 0 | 0 | 3 | 0 | 1 | 0 |
| E725K | 0 | 0 | 0 | 0 | 0 | 2 | 0 | T85I | 0 | 0 | 0 | 0 | 0 | 0 | 0 |
| G648R | 1 | 0 | 0 | 0 | 0 | 3 | 0 | F368V | 1 | 1 | 0 | 0 | 0 | 0 | 0 |
| G669R | 0 | 0 | 0 | 0 | 0 | 4 | 0 | G339S | 0 | 0 | 0 | 1 | 0 | 0 | 0 |
| L533K | 0 | 0 | 0 | 0 | 0 | 2 | 0 | P129S | 0 | 0 | 3 | 0 | 0 | 0 | 0 |
| L916F | 1 | 0 | 0 | 0 | 0 | 1 | 0 | E309A | 1 | 1 | 0 | 0 | 0 | 0 | 0 |
| P863H | 4 | 15 | 0 | 0 | 0 | 0 | 0 | A1914D | 2 | 3 | 0 | 0 | 0 | 0 | 0 |
| R905S | 2 | 0 | 0 | 0 | 0 | 0 | 0 | P1558L | 0 | 0 | 3 | 2 | 0 | 2 | 0 |
| S758I | 2 | 0 | 0 | 0 | 0 | 0 | 0 | L111S | 0 | 0 | 0 | 0 | 5 | 0 | 0 |
| S875F | 1 | 0 | 0 | 0 | 0 | 2 | 0 | G309C | 1 | 2 | 0 | 0 | 0 | 0 | 0 |
| T716P | 0 | 0 | 0 | 0 | 0 | 2 | 0 | P108S | 0 | 0 | 7 | 0 | 0 | 0 | 0 |
| T874P | 0 | 0 | 0 | 0 | 0 | 3 | 0 | I106S | 0 | 0 | 2 | 0 | 0 | 0 | 0 |
| V534G | 0 | 0 | 0 | 0 | 0 | 2 | 0 | L75F | 1 | 2 | 0 | 0 | 0 | 0 | 0 |
| Q57H | 12 | 342 | 7 | 0 | 0 | 7 | 0 | N142L | 0 | 5 | 0 | 0 | 0 | 0 | 0 |
| G251V | 0 | 0 | 0 | 0 | 0 | 5 | 4 | R279C | 0 | 2 | 0 | 0 | 0 | 0 | 0 |
| D155Y | 0 | 2 | 5 | 0 | 0 | 0 | 0 | G277S | 0 | 2 | 0 | 0 | 0 | 0 | 0 |
| G172C | 0 | 0 | 3 | 0 | 0 | 0 | 0 | H64N | 1 | 1 | 0 | 0 | 0 | 0 | 0 |
| G172V | 0 | 0 | 3 | 0 | 0 | 0 | 0 | D303G | 0 | 0 | 2 | 0 | 0 | 0 | 0 |
| N257D | 1 | 1 | 0 | 0 | 0 | 0 | 0 | T803I | 0 | 0 | 2 | 0 | 0 | 0 | 0 |
| W45R | 0 | 0 | 0 | 7 | 0 | 0 | 0 | M124I | 0 | 0 | 3 | 0 | 0 | 0 | 0 |
| E95K | 1 | 1 | 0 | 0 | 0 | 0 | 0 | P227S | 0 | 0 | 2 | 0 | 0 | 0 | 0 |
| P45L | 0 | 2 | 0 | 0 | 0 | 0 | 0 | S607I | 0 | 0 | 2 | 0 | 0 | 0 | 0 |
| A97V | 0 | 0 | 0 | 3 | 0 | 22 | 0 |  |  |  |  |  |  |  |  |

Table 6 gives us a microscopic view of the severity of the mutations that occurred in the dataset taken. In 82% of deleterious mutations protein stability has been decreased due to single point mutation. It is already observed that maximum mutations have occurred in the isolates which belong to clade GH. Out of 18 deleterious mutations happened in isolates from clade GH in 15 isolates (S194L, D936Y, P863H, W161L, F368V, E309A, A1914D, G309C, L75F, H64N, Q57H, N257D, E95K, P45L, V104F) stability have been decreased due to mutations. It can be observed at Table 7 that due to mutations majorly amino acid Glutamine and Serine are affected. Glutamine(Q) has been changed to Histidine(H), and Serine(S) changed to Leucine(L). The result may indicate that in Asian countries SARS-CoV-2 isolates responsible for COVID-19 majorly belong to the clades GR and GH. Among them mutations that occurred in isolates of clade GH are deleterious in nature, so have an impact on the biological function of proteins. The mutations also change the structural stability of proteins by making changes in free energy(Δ*G*).

**Table 6:** Investigating Stability of deleterious mutations in total dataset.

| Mutation type | Mutation | Stability | $\Delta\Delta G$ | Mutation type | Mutation | Stability | $\Delta\Delta G$ |
| --- | --- | --- | --- | --- | --- | --- | --- |
| S194L | Deleterious | Decreased | -0.47 | G77R | Deleterious | Decreased | -0.65 |
| D936Y | Deleterious | Decreased | -0.35 | T85I | Deleterious | Decreased | -0.91 |
| L752R | Deleterious | Decreased | -1.39 | F368V | Deleterious | Decreased | -1.74 |
| C851W | Deleterious | Decreased | -0.09 | G339S | Deleterious | Decreased | -1.2 |
| D820N | Deleterious | Decreased | -1.25 | P129S | Deleterious | Decreased | -1.82 |
| E725K | Deleterious | Decreased | -0.35 | E309A | Deleterious | Decreased | -0.96 |
| G648R | Deleterious | Decreased | -0.6 | A1914D | Deleterious | Decreased | -0.68 |
| G669R | Deleterious | Decreased | -0.14 | P1558L | Deleterious | Decreased | -0.36 |
| L533K | Deleterious | Decreased | -1.81 | L111S | Deleterious | Decreased | -2.29 |
| L916F | Deleterious | Decreased | -1.12 | G309C | Deleterious | Decreased | -1.07 |
| P863H | Deleterious | Decreased | -1.44 | P108S | Deleterious | Decreased | -1.73 |
| R905S | Deleterious | Decreased | -1.18 | I106S | Deleterious | Decreased | -2.24 |
| T716P | Deleterious | Decreased | -0.8 | L75F | Deleterious | Decreased | -1.09 |
| T874P | Deleterious | Decreased | -0.22 | R279C | Deleterious | Decreased | -0.67 |
| V534G | Deleterious | Decreased | -2.029 | G277S | Deleterious | Decreased | -1.3 |
| Q57H | Deleterious | Decreased | -0.9 | H64N | Deleterious | Decreased | -0.03 |
| G251V | Deleterious | Decreased | -0.54 | D303G | Deleterious | Decreased | -0.9 |
| G172C | Deleterious | Decreased | -0.83 | M124I | Deleterious | Decreased | -0.53 |
| G172V | Deleterious | Decreased | -0.41 | P227S | Deleterious | Decreased | -1.49 |
| N257D | Deleterious | Decreased | -0.21 | S193I | Deleterious | Increased | -0.29 |
| W45R | Deleterious | Decreased | -1.05 | R191L | Deleterious | Increased | -0.26 |
| E95K | Deleterious | Decreased | -0.58 | S180L | Deleterious | Increased | -0.12 |
| P45L | Deleterious | Decreased | -0.71 | A288P | Deleterious | Increased | -0.29 |
| V104F | Deleterious | Decreased | -1.47 | S758I | Deleterious | Increased | 0.43 |
| P38R | Deleterious | Decreased | -0.9 | S875F | Deleterious | Increased | 0.66 |
| W161L | Deleterious | Decreased | -0.51 | D155Y | Deleterious | Increased | 0.21 |
| R392C | Deleterious | Decreased | -1.29 | N142L | Deleterious | Increased | -0.05 |
| T113I | Deleterious | Decreased | -0.43 | T803I | Deleterious | Increased | 0.18 |
| M330T | Deleterious | Decreased | -1 | S607I | Deleterious | Increased | 0.37 |
| R287I | Deleterious | Decreased | -0.66 | A97V | Deleterious | Increased | -0.31 |
| P12S | Deleterious | Decreased | -1.29 |  |  |  |  |

**Table 7:** Amino Acid changed due to deleterious mutations taken place.

| MUTATION | COUNT | MUTATION | COUNT |
| --- | --- | --- | --- |
| Q>H | 368 | M>I | 3 |
| S>L | 215 | P>R | 3 |
| A>V | 25 | R>I | 3 |
| P>H | 19 | S>F | 3 |
| P>S | 16 | C>W | 2 |
| G>R | 12 | D>G | 2 |
| G>V | 12 | D>N | 2 |
| S>I | 12 | E>A | 2 |
| P>L | 9 | E>L | 2 |
| T>I | 9 | F>V | 2 |
| D>Y | 9 | H>N | 2 |
| W>R | 7 | I>S | 2 |
| G>C | 6 | L>K | 2 |
| A>D | 5 | L>R | 2 |
| L>F | 5 | L>S | 2 |
| N>L | 5 | M>T | 2 |
| T>P | 5 | N>D | 2 |
| E>K | 4 | R<sub>L</sub>>L | 2 |
| R>C | 4 | R>S | 2 |
| A>P | 3 | V>F | 2 |
| G>S | 3 | V>G | 2 |

## 4. Conclusions

The in silico analysis performed in this study states that the isolates in ten Asian countries are from clades G, GH, GR, L, S, O, and V. It indicates the diversity of the infection indeed. O, GH, and GR are the most widely affected ancestors of isolates among them. But when there is a talk about mutations, 31.93% of total mutations have taken place in the isolates of clade GR, and 31.33% of the mutations from GH. Hence, number of mutations are really high in the isolates belong to both the clades. When clades G, GH, and GR traversed almost in all countries specified here, the isolates of clade V are affected rarely. The most frequently mutated amino acids are Glutamine and Serine. In most of the cases glutamine is changed into Histidine and serine is changed to Leucine. It is to be noted that both the mutations are deleterious and the isolates of clade GH carry the major deleterious mutation load (44.19% of the total dataset). The majority of mutations taken place in the isolatess of clade GH are deleterious in nature. 82% of deleterious mutations are unstable and so their biological functions are affected. As a whole in this present work, the investigation provides us clade-wise characteristics of the SARS-CoV-2 isolates of the Asian continent. When reported research papers shed the light on development of clade-specific vaccines [35], our analysis can encourage drug designers for development of customized drugs or vaccines for Asian continent in order to combat COVID-19.

## Supporting information

*Table S1*. Country-wise specifications of isolates with the clades they belong to.

*Table S2*. Identifying mutations at different protein regions of the dataset taken.

## References

[1] R. Sanjuan, P. Domingo-Calap, Mechanisms of viral mutation, Cellular and molecular life sciences 73 (2016) 4433–4448.

[2] J. K. Das, A. Sengupta, P. P. Choudhury, S. Roy, Mapping sequence to feature vector using numerical representation of codons targeted to amino acids for alignment-free sequence analysis, Gene 766 (2020) 145096.

[3] D. Silvestro, A. Antonelli, N. Salamin, T. B. Quental, The role of clade competition in the diversification of north american canids, Proceedings of the National Academy of Sciences 112 (2015) 8684–8689.

[4] W. Tyor, C. Fritz-French, A. Nath, Effect of hiv clade differences on the onset and severity of hiv-associated neurocognitive disorders, Journal of neurovirology 19 (2013) 515–522.

[5] S. U. Kumar, D. T. Kumar, B. P. Christopher, C. Doss, The rise and impact of covid-19 in india, Frontiers in Medicine 7 (2020) 250.

[6] Y. Li, X. Yang, N. Wang, H. Wang, B. Yin, X. Yang, W. Jiang, The divergence between sars-cov-2 and ratg13 might be overestimated due to the extensive rna modification, Future Virology (2020).

[7] M. L. DeDiego, L. Pewe, E. Alvarez, M. T. Rejas, S. Perlman, L. Enjuanes, Pathogenicity of severe acute respiratory coronavirus deletion mutants in hace-2 transgenic mice, Virology 376 (2008) 379–389.

[8] K. G. Andersen, A. Rambaut, W. I. Lipkin, E. C. Holmes, R. F. Garry, The proximal origin of sars-cov-2, Nature medicine 26 (2020) 450–452.

[9] R. Zeng, R.-F. Yang, M.-D. Shi, M.-R. Jiang, Y.-H. Xie, H.-Q. Ruan, X.-S. Jiang, L. Shi, H. Zhou, L. Zhang, et al., Characterization of the 3a protein of sars-associated coronavirus in infected vero e6 cells and sars patients, Journal of molecular biology 341 (2004) 271–279.

[10] A. C. Walls, Y.-J. Park, M. A. Tortorici, A. Wall, A. T. McGuire, D. Veesler, Structure, function, and antigenicity of the sars-cov-2 spike glycoprotein, Cell (2020).

[11] A. Maitra, M. C. Sarkar, H. Raheja, N. K. Biswas, S. Chakraborti, A. K. Singh, S. Ghosh, S. Sarkar, S. Patra, R. K. Mondal, et al., Mutations in sars-cov-2 viral rna identified in eastern india: Possible implications for the ongoing outbreak in india and impact on viral structure and host susceptibility, Journal of Biosciences 45 (2020).

[12] N. K. Biswas, P. P. Majumder, Analysis of rna sequences of 3636 sars-cov-2 collected from 55 countries reveals selective sweep of one virus type, Indian J. Med. Res (2020).

[13] R. Kumar, H. Verma, N. Singhvi, U. Sood, V. Gupta, M. Singh, R. Kumari, P. Hira, S. Nagar, C. Talwar, et al., Comparative genomic analysis of rapidly evolving sars-cov-2 reveals mosaic pattern of phylogeographical distribution, Msystems 5 (2020).

[14] D. DiMaio, D. Nathans, Regulatory mutants of simian virus 40: effect of mutations at a t antigen binding site on dna replication and expression of viral genes, Journal of molecular biology 156 (1982) 531–548.

[15] A. Banerjee, R. Sarkar, S. Mitra, M. Lo, S. Dutta, M. Chawla-Sarkar, The novel coronavirus enigma: Phylogeny and analyses of coevolving mutations among the sars-cov-2 viruses circulating in india, JMIR Bioinformatics and Biotechnology 1 (2020) e20735.

[16] E. Foy, K. Li, C. Wang, R. Sumpter, M. Ikeda, S. M. Lemon, M. Gale, Regulation of interferon regulatory factor-3 by the hepatitis c virus serine protease, Science 300 (2003) 1145–1148.

[17] I. Astuti, et al., Severe acute respiratory syndrome coronavirus 2 (sars-cov-2): An overview of viral structure and host response, Diabetes & Metabolic Syndrome: Clinical Research & Reviews (2020).

[18] M. Eaaswarkhanth, A. Al Madhoun, F. Al-Mulla, Could the d614 g substitution in the sars-cov-2 spike (s) protein be associated with higher covid-19 mortality?, International Journal of Infectious Diseases (2020).

[19] S. S. Hassan, P. P. Choudhury, P. Basu, S. S. Jana, Molecular conservation and differential mutation on orf3a gene in indian sars-cov2 genomes, Genomics (2020).

[20] F. Pereira, Evolutionary dynamics of the sars-cov-2 orf8 accessory gene, Infection, Genetics and Evolution 85 (2020) 104525.

[21] S. M. Lokman, M. Rasheduzzaman, A. Salauddin, R. Barua, A. Y. Tanzina, M. H. Rumi, M. I. Hossain, A. Z. Siddiki, A. Mannan, M. M. Hasan, Exploring the genomic and proteomic variations of sars-cov-2 spike glycoprotein: a computational biology approach, Infection, Genetics and Evolution (2020) 104389.

[22] B. K. Kumar, A. Rohit, K. S. Prithvisagar, P. Rai, I. Karunasagar, I. Karunasagar, Deletion in the c-terminal region of the envelope glycoprotein in some of the indian sars-cov-2 genome, Virus Research (2020) 198222.

[23] R. Dutta, L. Buragohain, P. Borah, Analysis of codon usage of severe acute respiratory syndrome corona virus 2 (sars-cov-2) and its adaptability in dog, Virus research 288 (2020) 198113.

[24] I. Saha, N. Ghosh, D. Maity, N. Sharma, K. Mitra, Inferring the genetic variability in indian sars-cov-2 genomes using consensus of multiple sequence alignment techniques, Infection, Genetics and Evolution 85 (2020) 104522.

[25] S. S. Hassan, P. P. Choudhury, B. Roy, S. S. Jana, Missense mutations in sars-cov2 genomes from indian patients (2020).

[26] S. S. Hassan, P. P. Choudhury, B. Roy, Sars-cov2 envelope protein: non-synonymous mutations and its consequences (2020).

[27] J.-Y. Li, C.-H. Liao, Q. Wang, Y.-J. Tan, R. Luo, Y. Qiu, X.-Y. Ge, The orf6, orf8 and nucleocapsid proteins of sars-cov-2 inhibit type i interferon signaling pathway, Virus research 286 (2020) 198074.

[28] S. Weber, C. Ramirez, W. Doerfler, Signal hotspot mutations in sars-cov-2 genomes evolve as the virus spreads and actively replicates in different parts of the world, Virus research 289 (2020) 198170.

[29] L. Li, T. Sun, Y. He, W. Li, Y. Fan, J. Zhang, Epitope-based peptide vaccines predicted against novel coronavirus disease caused by sars-cov-2, BioRxiv (2020).

[30] Q. Nie, X. Li, W. Chen, D. Liu, Y. Chen, H. Li, D. Li, M. Tian, W. Tan, J. Zai, Phylogenetic and phylodynamic analyses of sars-cov-2, Virus research 287 (2020) 198098.

[31] P. Priya, A. Shanker, Coevolutionary forces shaping the fitness of sars-cov-2 spike glycoprotein against human receptor ace2, Infection, Genetics and Evolution (2020) 104646.

[32] M. B. Uddin, M. Hasan, A. Harun-Al-Rashid, M. I. Ahsan, M. A. S. Imran, S. S. U. Ahmed, Ancestral origin, antigenic resemblance and epidemiological insights of novel coronavirus (sars-cov-2): Global burden and bangladesh perspective, Infection, Genetics and Evolution 84 (2020) 104440.

[33] M. Seyran, D. Pizzol, P. Adadi, T. M. A. El-Aziz, S. S. Hassan, A. Soares, R. Kandimalla, K. Lundstrom, M. Tambuwala, A. A. Aljabali, et al., Questions concerning the proximal origin of sars-cov-2, Journal of Medical Virology (2020).

[34] D. Mercatelli, F. M. Giorgi, Geographic and genomic distribution of sars-cov-2 mutations (2020).

[35] C. Yi, X. Sun, J. Ye, L. Ding, M. Liu, Z. Yang, X. Lu, Y. Zhang, L. Ma, W. Gu, et al., Key residues of the receptor binding motif in the spike protein of sars-cov-2 that interact with ace2 and neutralizing antibodies, Cellular & Molecular Immunology (2020) 1–10.

[36] K. Boonnak, Y. Matsuoka, W. Wang, A. L. Suguitan, Z. Chen, M. Paskel, M. Baz, I. Moore, H. Jin, K. Subbarao, Development of clade-specific and broadly reactive live attenuated influenza virus vaccines against rapidly evolving h5 subtype viruses, Journal of Virology 91 (2017).

[37] Y. Choi, A. P. Chan, Provean web server: a tool to predict the functional effect of amino acid substitutions and indels, Bioinformatics 31 (2015) 2745–2747.

[38] E. Capriotti, P. Fariselli, R. Casadio, I-mutant2. 0: predicting stability changes upon mutation from the protein sequence or structure, Nucleic acids research 33 (2005) W306–W310.

[39] Y. Choi, G. E. Sims, S. Murphy, J. R. Miller, A. P. Chan, Predicting the functional effect of amino acid substitutions and indels, PloS one 7 (2012) e46688.

[40] E. Capriotti, P. Fariselli, I. Rossi, R. Casadio, A three-state prediction of single point mutations on protein stability changes, BMC bioinformatics 9 (2008) S6.

[41] S. Chun, J. C. Fay, Identification of deleterious mutations within three human genomes, Genome research 19 (2009) 1553–1561.

